# Effect of artificial light at night on sleep and metabolism in weaver birds

**DOI:** 10.1101/2021.07.03.451020

**Authors:** Anupama Yadav, Raj Kumar, Jyoti Tiwari, Vaibhav Vaish, Shalie Malik, Sangeeta Rani

## Abstract

Artificial light at night is constantly minimizing the span of dark nights’ from the natural light-dark cycle of earth. Over the past century, the ‘lightscape’ of earth has completely changed owing to technological advancements which subsequently changed the lifestyle of human as well as the nearby animal species. This motivated the present study, wherein we investigated the impact of light at night (LAN) on behavior and physiology of a diurnal passerine finch, baya weaver (*Ploceus philippinus*). A group of bird (N=10) exposed to 12L:12D photoperiod was initially subjected to dark nights (0 lux) for a period of 10 days followed by 5 lux; night light for a span of 4 weeks. First week in LAN served as acute treatment with respect to fourth week (chronic). Acute exposure had more pronounced impact on the behavioral and physiological observations when compared with chronic treatment. The results reveal significant increase in nighttime activity, sleep loss, significant inclusion of drowsiness behavior during the day in response to LAN. Beside these behavioral alterations, changes in physiological parameters such as; reduction in body mass, loss of gradient between pre- and post-prandial blood glucose levels, elevation in plasma corticosterone levels were more prominent during acute exposure of LAN. Plasma metabolites such as triglycerides, total protein, serum glutamic-oxaloacetic transaminase (SGOT) and creatinine concentrations also hiked in response to LAN treatment. Thus, the study broadly enumerates the impact of acute and chronic exposure of LAN on behavior (rest/sleep) and physiology (metabolism) of birds’.

## 1. Introduction

Life on earth has evolved around the diel and seasonal variation of light – dark cycle. Since millions of years, organisms have internalized the light – dark system to optimize the resource utilization in their spatial niches and minimize survival risks (Russart and Nelson, 2018). Over the past century, the ‘lightscape’ of earth has completely changed owing to the expansion of human habitation near and within natural habitats. The expanse of LAN has been increasing continuously across the globe since the advent of electricity (Kyba et al. 2017). LAN affects organisms’ both directly (via lit infrastructure from urban setting) and indirectly (via sky glow i.e. reflections of urban illumination by cloud cover or air borne particulate matter) (Bennie et al. 2015, 2016).

Studies on LAN report an altered orientation of basic response and function in time and space. Since time immemorial, photoperiod is considered as a proximate cue regulating different circadian and seasonal attributes of a living system (Kumar et al. 1997). Organisms’ have developed time measurement system i.e. circadian clocks’ which allows them to entrain and align different behavioral and physiological activities to appropriate time of day (Falcon et al. 2020). For instance, daily variations of sleep, locomotor activity, food intake, body temperature, hormone secretion and gene expression, all are fine tuned by the light-dark cycle (Foster and Kreitzmann 2004, Roenneberg et al. 2007, Wright et al. 2013, Azzi et al. 2014, Welbers et al. 2017). Studies from the past report the deleterious effects of LAN on animals ranging from invertebrates to vertebrates (Falcon et al. 2020). Amongst vertebrates, birds are the most explored model system with respect to studies related to LAN (Falcon et al. 2020). The probable reason for higher usage of birds for these experiments may be their closeness to human habitation, besides this, they offer two variants as resident and migratory which makes them more potent experimental model that can reveal information related to local disturbances along with latitudinal impacts of LAN on birds’ behavior and physiology. Studies on both resident and migratory species reveal the detrimental impact of night light on their circadian and circannual biology. LAN attracts on-route migratory birds which might result in either collision, leading to fatal consequences (Ronconi et al. 2015, Rodriguez et al. 2017) or disorientation of migratory track (Van Doren et al. 2017). In case of path diversion, migratory birds end up losing both time and energy which ultimately influences their survival. Studies on resident species, subjected to visible spectrum have reported the impact of LAN on several daily rhythms such as locomotor activity, sleep, bird song, body temperature, melatonin secretion, immunity and oxidative stress in Great Tit Parus major (Ouyang et al., 2015; Raap et al., 2015, 2016a,c; de Jong et al., 2016a, 2017; Raap, 2018), Blackbird Turdus merula (Dominoni et al., 2013b), Indian Weaver Bird Ploceus philippinus (Singh et al., 2012, Kumar et al., 2018), Zebra Finches Taeniopygia guttata (Moaraf et al., 2020). Besides affecting the circadian biology, LAN significantly affects the annual reproductive decision in birds. In Blackbird Turdus merula, LAN phase advanced the annual cycle of gonadal maturation, steroid levels and body molt (Dominoni et al., 2013b). Similar observations were obtained from California Jay, Aphelocoma californica, wherein the plasma testosterone, oestradiol, melatonin and LH levels showed sex-specific alterations under the influence of night light (Schoech et al., 2013). In Mockingbirds Mimus polyglottos and American Blackbirds T. migratorius, LAN altered the onset of dawn song and courtship behavior during breeding season in a dose-dependent manner (Longcore, 2010). Since, these clock dependent functions are tightly coupled with natural light-dark cycles orchestrating myriad of downstream events and anthropogenic night light has apparently changed the perception of the daylength, this results in a mismatch between an organisms’ endogenous rhythm and the external environment i.e. ‘circadian desynchrony’ (Wyse et al. 2011). This circadian desynchrony may lead to altered behavior and physiology which subsequently affects the health and fitness of organisms’.

Most of the studies discussed above generally report the short term impact of LAN exposure. Chronic effects of night light are also reported in some studies, but a comparative account of acute and chronic exposure of LAN suggesting an adaptive approach of the organism’s when dealing with LAN is missing to best of our knowledge. Therefore, in the present study we have tried to investigate the acute and chronic response of LAN on behavior and physiology of baya weaver (*Ploceus philippinus*) bird. This is a diurnal, photoperiodic, long day-breeder species, spanning across Indian subcontinent and Southeast Asia. In order to gain better understanding of night light related adjustments, we have assessed different behavioral attributes of the birds such as distribution of locomotor activity, changes in activity-rest pattern, and alterations in sleep behavior with special reference to different sleep postures. Besides these, different physiological factors were also considered for the study, for instance, body mass, body temperature rhythm and measurement of different plasma metabolites such as total protein, triglyceride, creatinine, uric acid, bilirubin total and serum glutamic oxaloacetic transaminase (SGOT), blood glucose and plasma corticosterone levels were assessed to evaluate the illuminated nights’ stress. Thus, besides the discrete response of acute and chronic treatment of LAN, we were interested in understanding if the birds can adapt to prolong exposure of light at night?

## 2. Methods

### 2.1. Birds and maintenance

The experiment was conducted on adult male weaver birds captured from nature by mist netting. Thereafter, they were brought to laboratory where they were kept in an outdoor aviary (size = 30.0 × 2.5 × 2.5m) for 2 weeks, for acclimation. The outdoor aviary offered natural daylength and temperature conditions besides, it was supplemented with plants and grasses to provide a natural ambience to the birds. During acclimation food and water was provided ad libitum and replenished on daily basis during the daytime. Food mainly constituted of seeds of *Oryza sativa*. Besides the seeds, birds were also provided with a customized supplement food rich in protein and vitamins twice a week to maintain them in healthy condition prior to experiment. After 2 weeks, birds were brought indoors and kept in 12L:12D (12 hours of light and 12 hours of darkness) photoperiodic condition before they were subjected to the experimental schedule.

### 2.2. Experimental protocol

The experiment was designed to assess the impact of acute and chronic exposure of LAN on behavior and physiology of birds. In a longitudinal study, the weaver birds (N = 10) were subjected to dark nights (nighttime intensity = 0 lux) followed by night light (5 lux intensity), for an experimental schedule of about 6 weeks. After initial acclimation of 3-4 days in the activity recording chamber, the subsequent week with dark nights served as control during the experiment. Afterwards, the birds were exposed to lit up nights (5 lux) for 4 consecutive weeks. First week of LAN exposure was considered as acute treatment whereas the response at the end of 4^th^ week served as chronic. The experiment was carried out in a light and temperature regulated facility. Birds were subjected to 12L:12D photoperiodic schedule at 24 ± 2 □ C ambient temperature during entire experiment. Birds were singly housed in activity recording chamber (size = 60 × 45 × 35 cm) supplemented with two light sources (compact fluorescent lamps; CFL of 11W and 5W, 230V, Phillips, India). The two lamps were used alternatively during day and night respectively with the desired radiance output. During the day the birds were subjected to 110 ± 2 lux luminance, whereas, during night, 5 ± 00.05 lux intensity was set at perch level, as measured by Q203 Quantum Radiometer (Macam Photometrix Ltd., Scotland, UK). The desired illumination was obtained with the help of perforated black paper hood placed over the light source. Sampling for different behavioral and physiological parameters was done weekly during control, acute and chronic treatment of LAN.

Various parameters assessed in this study and their methodology is summarized as follows;

### 2.3. Activity-rest pattern

The locomotor activity was recorded on daily basis with the help of infrared motion sensors (Haustier PI-meldor; C and K systems, Conrad Electronics, Hirschau, Germany) mounted on each recording chamber, as mentioned above. The movement of birds’ inside their cage was recorded in 5 minute bin size, over a period of 24 hour whose output was recorded and stored as separate file in a computerized system. The Chronobiology Kit program (Stanford Software System, USA) was used for retrieving these activity records (actogram) and for further calculation and analysis of total counts, activity-rest duration and daily onset and offset of activity. Weekly data of activity during each treatment phase namely; control (no LAN), acute (1^st^ week LAN) and chronic (4^th^ week LAN) was considered for analysis. During calculations when in a span of 5 minutes no activity bouts were present, that particular duration was considered as rest. The calculation has been processed on hourly basis and was continued for duration of 24 hour up till 7 days of week.

### 2.4. Sleep behavior

Sleep behavior was recorded following Yadav et al. (2021). Briefly, birds were monitored for different active and sleep behaviors during a 24 hour day. Active behaviors included active and alert wakefulness whereas sleep was assessed by front sleep, back sleep and uni-hemispheric sleep (UHS) postures (Fuchs et al. 2006, Yadav et al. 2017). Drowsiness was considered as an intermediate state between sleep and wake, thus omitted from calculations for total sleep but was used for calculating rest. Sleep was recorded equally between day and night having three time points in each phase of day followed by two transitions i.e. from day to night and vice-versa. In all sampling was done at 8 time points which were an hour long: i) midday (MD) / midnight (MN) ii) earlyday (ED) / early night (EN) iii) lateday (LD) / late night (LN) and iv) transition from day to night and vice-versa (trans1 and trans 2). Thus, sleep was recorded at 8 equispaced points during a 24 hour day for two consecutive days during different treatment schedules in the experiment.

### 2.5. Food intake

Food intake was measured following Jain and Kumar (1995). A known quantity of food was given to the birds, leaving it undisturbed for two successive days. At the end of two-cycles of 24 hour each, the remaining food was removed carefully from the cage and was measured after removing the faecal matter. The difference between food supplied and food left gave the amount of food consumed (food intake) by the birds.

### 2.6. Body mass and body temperature

Body mass and body temperature were measured at the end of the week during control and acute treatment, whereas at the end of 4^th^ week during chronic treatment in the experiment. Body mass was measured using a top pan balance with an accuracy of 00.01 gram (Schimadzu ELB 300). Briefly the birds were wrapped in a cotton bag and weighed just after lights on in the morning. Body temperature was measured using infrared Thermoscan (Model = EXP-01B) from the flank area of the birds (Kumar et al. 2018). The surface temperature was also recorded for two consecutive cycles of 24 hour each at 6 time points (Zeitgeber time; ZT-02, 06, 10, 14, 18 and 22) of the day.

### 2.7. Biochemical Parameters

The observations for different biochemical parameters from blood plasma were done at middle of the day (ZT 06) and night (ZT 18) to assess the impact of acute and chronic exposure of LAN on bird’s physiology. Blood sample was obtained by puncturing the left brachial vein of the birds. For harvesting the plasma, 100-150 µl blood was collected in a heparinised capillary tube and thereafter, transferred to microcentrifuge tube. Afterwards the blood was centrifuged for 15 minutes at 4 □ C at 3000 r.p.m. to harvest the plasma. The plasma retrieved was stored at -80 □ C until processed for the different assays.

Plasma corticosterone levels were measured by using immunoassay kit from Enzo Life Sciences (catalog no. ADI-900-097) during the day (ZT 06) and night (ZT 18) following manufacturers’ protocol (Crino et al., 2014; Mishra et al., 2017). Briefly, the assay was performed by using 10 μl of plasma sample in 1:40 dilution. Initially 100 μl of standards and plasma samples were placed in the standard and sample wells respectively followed by 100 μl of assay buffer in NSB and blank (B_0_) wells. Then 50 μl of assay buffer was added to NSB wells and 50 μl blue conjugate (alkaline phosphate conjugated with cort) to each well, except B_0_ and TA (total activity) wells. Subsequently 50 μl of antibody was added to each well, except NSB, B_0_ and TA wells. Thereafter the plate was incubated on a shaker at 500 rpm for 2 h at room temperature. Following 3 washes with wash-buffer, 5 μl conjugate was added to TA wells and 200 μl of p-nitrophenyl phosphate in buffer (pNpp) to every well and the plate was kept at room temperature without shaking for 1 h. At the end, 50 μl of stop-solution was added to stop the reaction, and the plate was read at 405 nm by *Biorad 680 microplate reader (USA)*. The sensitivity and intra-assay variability of the assay were 26.99 pg/ml and 8.4%, respectively.

Biochemical assay kits from Erba diagnostics Mannheim GmbH, Germany, were used to analyze bilirubin total, creatinine, triglycerides, total protein, uric acid and SGOT (Serum glutamic-oxaloacetic transaminase) from plasma samples of birds in different LAN treatment phases. The kit is widely used in assessing the different biochemical parameters in mammalian and avian model systems (Shivprasad et al., 2014, Behera et al., 2019, Khandia et al., 2020). All samples were diluted in the ratio of 1:3 (plasma: distilled water) to attain sufficient volumes for running all the tests. All the kits were run according to the manufacturer’s protocol to obtain the results. The biochemical parameters were analyzed through *Csense 100 Auto Chemistry Analyzer* (*Medsource Ozone Biomedicals Private Limited, New Delhi, India*).

Levels of blood glucose were measured at pre-prandial (ZT 22) and post-prandial (ZT 02) time points with the help of Accu check active from Roche Diagnostics (GmbH Mannheim, Germany) by placing a drop of blood (∼5µl) on *Accu chek active strip* (Lieske et al., 2002, Singh et al., 2016).

### 2.8. Statistics

All the statistical analyses were performed on GraphPad Prism version 5. Most of the behavioral and physiological datasets were analyzed using 1-way or 2-way repeated measure analysis of variance (RM ANOVA) as there were broadly three experimental conditions (control, acute LAN and chronic LAN) followed by Newman-keuls or Bonferroni post hoc tests respectively for multiple comparisons when ANOVA gave significant results. 2-way RM ANOVA was deployed to test the parameters subjected to different exposure of LAN (factor 1) along with their response at different times of day (factor 2). Besides, the usage of RM ANOVA, two-tailed-t-test was used to calculate the significance while analyzing the change in body mass and food intake data. Furthermore, unimodal cosinor regression analysis (y = A +[B.cos (2π (x-C)/24)]), where A, B and C denoted the mean (mesor), amplitude and acrophase of the rhythm, respectively (Cuesta et al. 2009) was used to evaluate the presence and persistence of rhythmic character of different sleep behavior studied during the course of experiment. The significance of cosinor analysis was calculated with the help of Daniel Soper, an online software (http://www.danielsoper.-com/statcalc3/calc.aspx?id=15, Soper 2013).

## 3. Results

### 3.1. Distribution of locomotor activity and activity-rest pattern

The representative actogram clearly reflects the impact of LAN on locomotor activity pattern and its distribution (Fig.1). Weaver birds being diurnal showed only daytime activity as was evident from the control phase (dark nights) of treatment, the activity can be seen restricted during the day (12L) whereas, the night (12D) was mostly clean with minor bouts of activity. With the exposure of LAN, there was significant increase in the nighttime activity. The 24 hour activity profile of the birds under different treatment phases clearly reflects the intensity of disturbance in the night during acute exposure (P = 0.0001, F_2,46_ = 11.11; 1-way RM ANOVA, Fig.1a). Continuous exposure of LAN has gradually decreased the intensity of disturbance but it was not completely abolished (P = 0.0001, F_2,46_ = 11.11; 1-way RM ANOVA, Fig.1a). Taking into account the daytime (P = 0.0300, F_2,18_ = 4.290), nighttime (P = 0.0008, F_2,18_ = 10.82) and a sum total of activity throughout the day (P = 0.0278, F_2,18_ = 4.398), significant impact of lit-up nights was visible when compared with dark night (1-way RM ANOVA, Fig.1b). Daytime activity was solely responsible for all the activity bouts of birds in control phase. Acute exposure of LAN had approximately similar day counts as that of control but a significant higher proportion of nighttime activity resulted in overall increase in total activity counts of the birds. Chronic exposure of LAN also had significant proportion of night counts in comparison with control, but the daytime activity was compromised which resulted in gross reduction of total counts of birds across the day. An appreciable impact of lit up nights was also observed on the onset and offset of activity of birds. Exposure of LAN has altered the phase relation between lights on and off and activity onset and offset respectively. Activity onset has significantly advanced with initial exposure of LAN, and was seemingly fixed, as the difference between activity onset and lights-on was nearly constant with time (P <0.0001, F_4,36_ = 20.76, 1-way RM ANOVA, Fig.1c). Contrarily, activity offset significantly delayed, followed by, gradual decrease in the difference between activity offset and lights-off with passage of time (P <0.0001, F_4,36_ = 11.17, 1-way RM ANOVA, Fig.1c).

**Figure 1:**
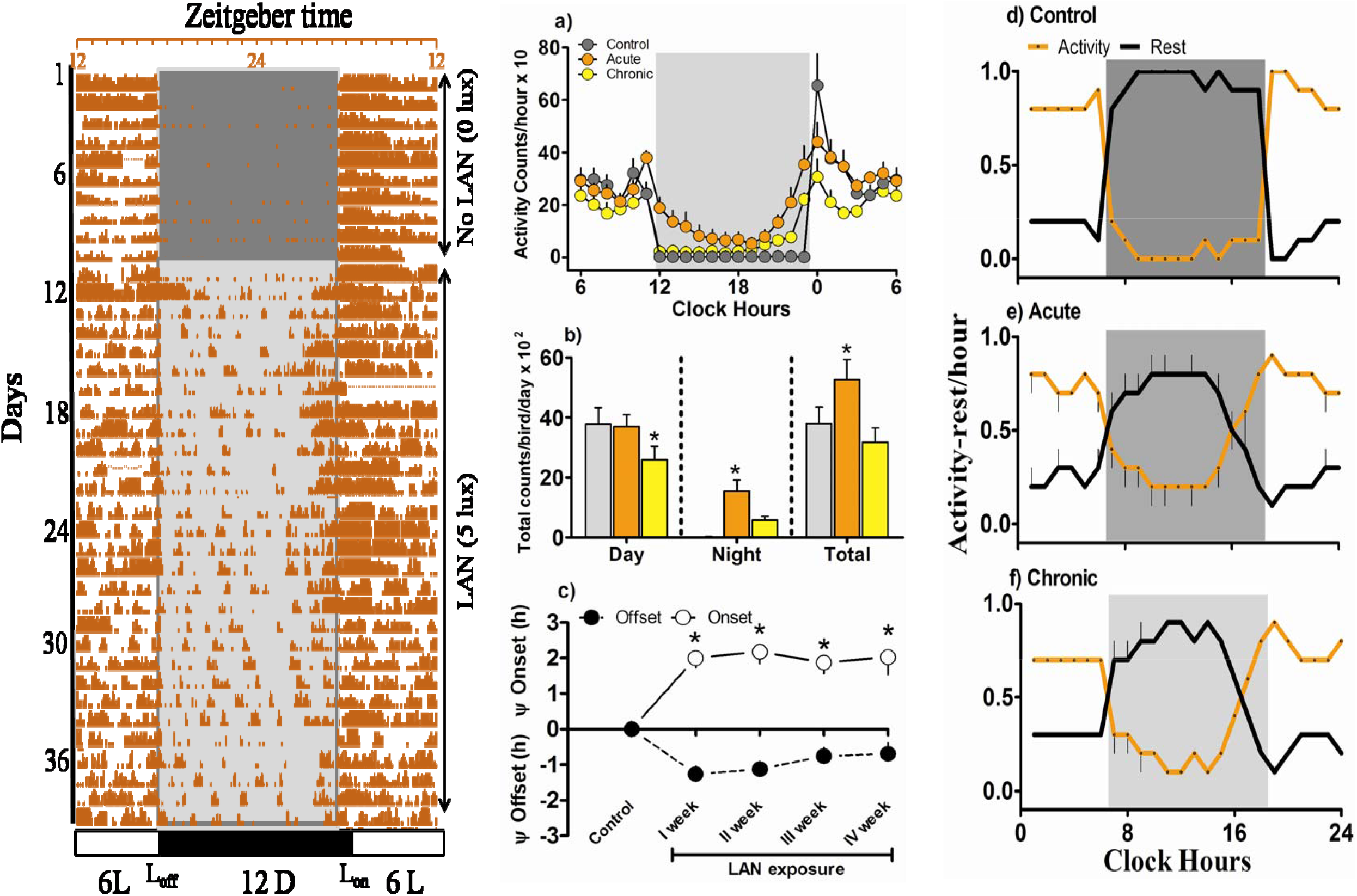
Distribution of activity-rest across the day in baya weaver birds exposed to LAN. Representative actogram of baya weaver bird subjected to dark nights (0 lux) followed by night light of 5 lux (left panel). Mean ±SEM values of ; 24 hour profile of activity counts of birds in the three treatment phases, namely control, acute and chronic LAN (a), Diel distribution of total activity bouts of birds in different treatment phases (b), weekly distribution of onset and offset of activity during the course of experiment (c). Graphs (d-f) represent mean ±SEM values of activity and rest in control, acute and chronic LAN treatment respectively. Asterisk (*) above the bar represent the significant difference between the phases. Significance was considered at p < 00.05.

Distribution of activity and rest followed nearly the same pattern for control and chronic treatment during the day, whereas, acute and chronic treatments’ resembled with each other during course of night. In the absence of night-light, the initial four hours of the day beginning with lights-on, contributed maximally to the active phase but the same does not stands true for chronic exposure of LAN. Generally, the birds can be observed resting during second half of the day in control and chronic treatment (Fig.1d, f), but acute exposure of LAN has fragmented this resting regime of birds (Fig.1e). No crossover between activity and rest can be visualized in the dark phase during absence of night-light, but since exposure of LAN advanced the activity onset, this resulted in intersection between activity and rest in the last quarter of night. The results clearly represent high activity duration (interaction; P <0.0001, F_2,36_ = 31.31, treatment; P = 0.0034, F_2,36_ = 6.701, time of day; P <0.0001, F_1,36_ = 209.7, 2-way RM ANOVA) and compromised rest period (interaction; P <0.0001, F_2,36_ = 31.23, treatment; P = 0.0030, F_2,36_ = 6.866, time of day; P <0.0001, F_1,36_ = 208.4, 2-way RM ANOVA) during night because of artificial light at night.

### 3.2. Sleep behavior and postural reallocation

Sleep was quantified with the help of different behavioral postures as illustrated in the stack plot (Fig.2). Awake state was represented by active and alert wakefulness, whereas sleep was assessed via, front sleep, back sleep and uni-hemispheric sleep (UHS) behaviorally. Drowsiness acted as an intermediate state linking sleep and wakefulness, thus was categorized as resting zone behavior rather than a typical sleep state. Talking with respect to different treatment phases, during control phase (dark nights, Fig.2a), the day was marked by significant presence of active and alert behaviors with traces of UHS and drowsiness. The transition from day to night had abundance of active wakefulness followed by alert wakefulness during the day component of transition, whereas the night component constituted maximally of alert wakefulness followed by drowsiness and front sleep with traces of active wakefulness. Different phases of night, namely, early night, mid night and late night constituted of substantial proportions of front sleep and back sleep. Front sleep can be observed being overtaken by back sleep with progression of night. Drowsiness and alert wakefulness had minor contribution during early and mid night phases which subsequently abolished in the late night phase. The night component of the second transition from dark to light, constituted majorly of different rest/sleep behaviors, with minor proportions of alert wakefulness whereas the day component comprised only of active and alert wakefulness.

**Figure 2:**
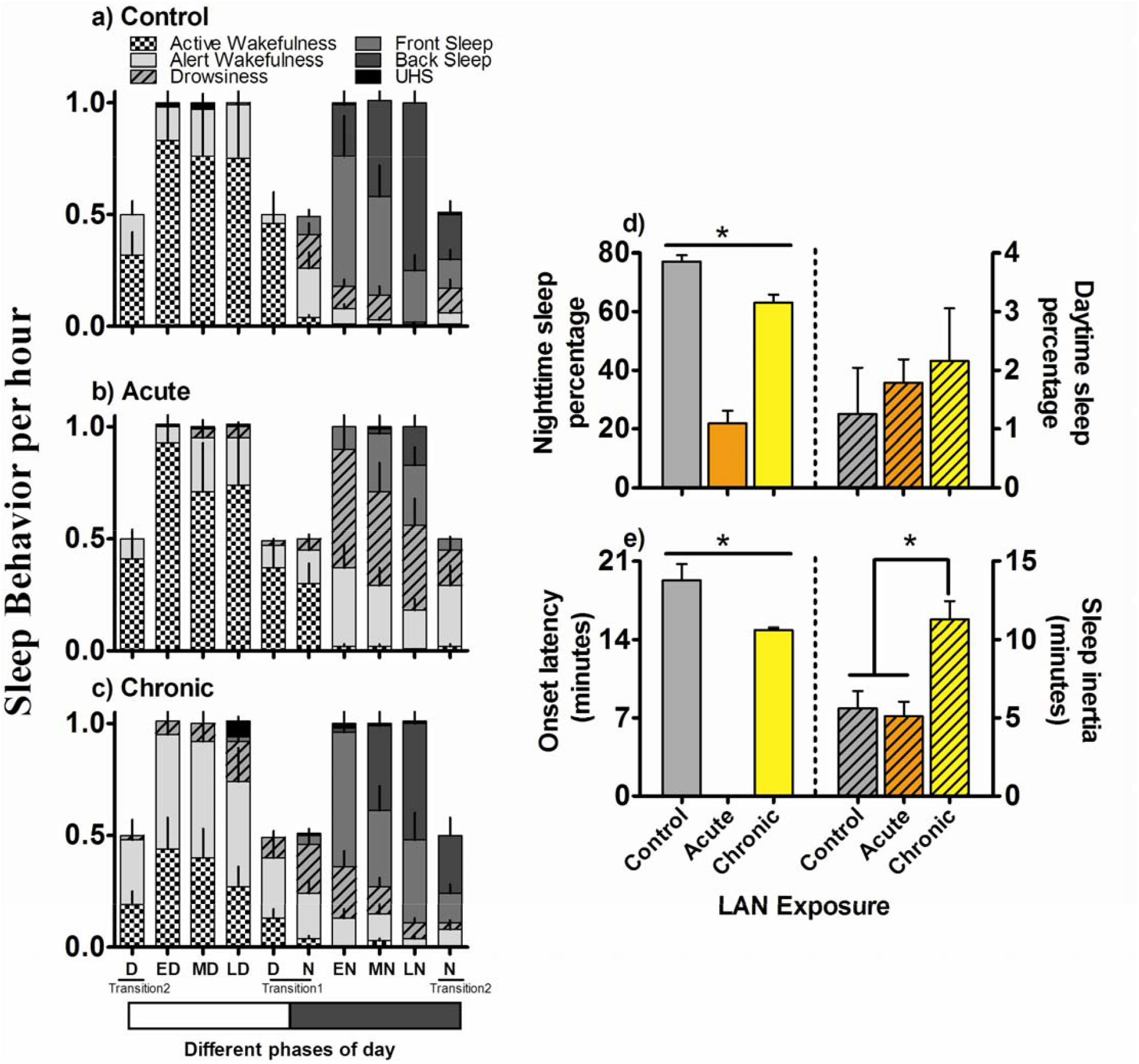
Diel distribution of different sleep behaviors and sleep characteristics of baya weaver when exposed to LAN. Stack plot of different sleep behaviors presented as mean ±SEM values spread across the day and at different LAN treatment schedules (graphs a-c). Sleep percentage of birds in different treatment phases during day and night (d). Mean ±SEM values of sleep onset latency and sleep inertia in birds (e). Asterisk (*) above the bar represent the significant difference between the phases. Significance was considered at p < 00.05.

Acute exposure of LAN dramatically affected the different night phase behaviors (Fig.2b). As evident from the graph, now the night can be seen dominated by drowsiness. Early phase of night constituted majorly of drowsiness and alert wakefulness with small portion of front sleep depicting the impact of lit-up nights. With subsequent progression of night, amount of front and back sleep has increased but still the dominance of drowsiness and alert wakefulness can be visualized from the stack plot. Currently the night component of transitions’ were also significantly marked by the abundance of active and alert wakefulness followed by drowsiness and minor bouts of front sleep (specifically in the transition from dark to light). The daytime behaviors were not much affected by the night light, but now minor incorporation of drowsiness was observed in second half of the day as well. The day component of transition had the same behavioral composition as that of control phase with majority of active wakefulness followed by alert wakefulness.

Chronic exposure (Fig.2c) of LAN resulted in relapse of nighttime sleep behaviors when compared with acute exposure. Now the early night phase showed generous proportion of front sleep followed by drowsiness and alert wakefulness. The proportion of front and back sleep increased gradually with the passage of time but there was no abolition of drowsiness and alert wakefulness in any phase of night. The night part of both the transitions’ showed resemblance with its counterpart in control phase (dark night) treatment. Interestingly chronic exposure resulted in addition of drowsiness to every component of day. Besides this, the later half of the day also reported the presence of back and front sleep postures reflecting the reallocation of different sleep and resting behaviors because of night light. Table 1 summarizes the results of two-way RM ANOVA supporting the aforementioned description (duration of LAN treatment served as, factor 1 and time of day as, factor 2).

**Table 1.** Results of 2 way-RM ANOVA

| **Active Behavior** | **Factors** | **Df** | **F value** | **P value** |
| --- | --- | --- | --- | --- |
| Active Wakefulness | Interaction | 18 | 2.618 | **0.0012** |
|  | LAN treatment | 2 | 5.118 | **0.0247** |
|  | Time of Day | 9 | 46.41 | **<0.0001** |
| Alert Wakefulness | Interaction | 18 | 2.519 | **0.0018** |
|  | LAN treatment | 2 | 2.443 | **0.0128** |
|  | Time of Day | 9 | 3.202 | **0.0018** |
| **Sleep Behavior** |  |  |  |  |
| Drowsiness | Interaction | 18 | 5.822 | **<0.0001** |
|  | LAN treatment | 2 | 15.77 | **0.0004** |
|  | Time of Day | 9 | 11.78 | **<0.0001** |
| Front Sleep | Interaction | 18 | 2.311 | **0.0043** |
|  | LAN treatment | 2 | 4.623 | **0.0325** |
|  | Time of Day | 9 | 23.00 | **<0.0001** |
| Back Sleep | Interaction | 18 | 3.729 | **<0.0001** |
|  | LAN treatment | 2 | 8.280 | **0.0055** |
|  | Time of Day | 9 | 23.24 | **<0.0001** |
| UHS | Interaction | 18 | 1.048 | 0.4145 |
|  | LAN treatment | 2 | 0.768 | 0.4852 |
|  | Time of Day | 9 | 1.141 | 0.3405 |

Result of 2 way-RM ANOVA of different sleep and active behaviors of baya weaver across different points of the day during observation period. The significant test responses ( $p < 0.05$ ) are reported in bold in the table.

Calculation of sleep percentage revealed significant reduction in nighttime sleep during acute exposure of LAN (interaction; P <0.0001, F_2,36_ = 70.37, treatment; P < 0.0001, F_2,36_ = 68.82, time of day; P <0.0001, F_1,36_ = 917.3; 2-way RM ANOVA) whereas the daytime sleep seemed to be unaffected by LAN (Fig.2d). Sleep onset latency and sleep inertia also varied significantly in response to LAN (interaction; P <0.0001, F_2,36_ = 52.66, treatment; P < 0.0001, F_2,36_ = 80.62, time of day; P = 0.0001, F_1,36_ = 24.33; 2-way RM ANOVA). As observed in Fig. (2e) no onset of sleep could be observed in acute exposure of LAN during the observation period. Chronic exposure of LAN resulted in increased sleep inertia in birds after lights on (P < 0.001; Bonferroni post test).

Besides, the distribution and allocation of different sleep and awake behaviors, amplitude and rhythmicity of different behaviors across 24 hour was also assessed (supplementary Fig.1). All the behaviors mentioned during the course of study showed rhythmic pattern across the day except for the amplitude which changed dramatically with the introduction of night light. The amplitude and rhythmicity of different sleep behaviors namely; front, back and uni-hemispheric sleep was approximately similar for control and chronic treatment of LAN, whereas in case of acute exposure, the aforementioned characters of these behaviors had nearly flattened. Drowsiness showed its surge during nighttime in acute treatment whereas in case of chronic exposure its amplitude significantly increased for every phase of day. Active wakeful state resembled each other in amplitude and oscillation during control and acute treatment whereas there was significant reduction in amplitude of this behavior during chronic exposure of LAN. Contrarily, alert wakefulness increased significantly in amplitude during chronic exposure in comparison to control and acute treatment phase. Table 2 summarizes the results of cosinor analysis revealing the time of maxima of each behavior analyzed.

**Table 2.**
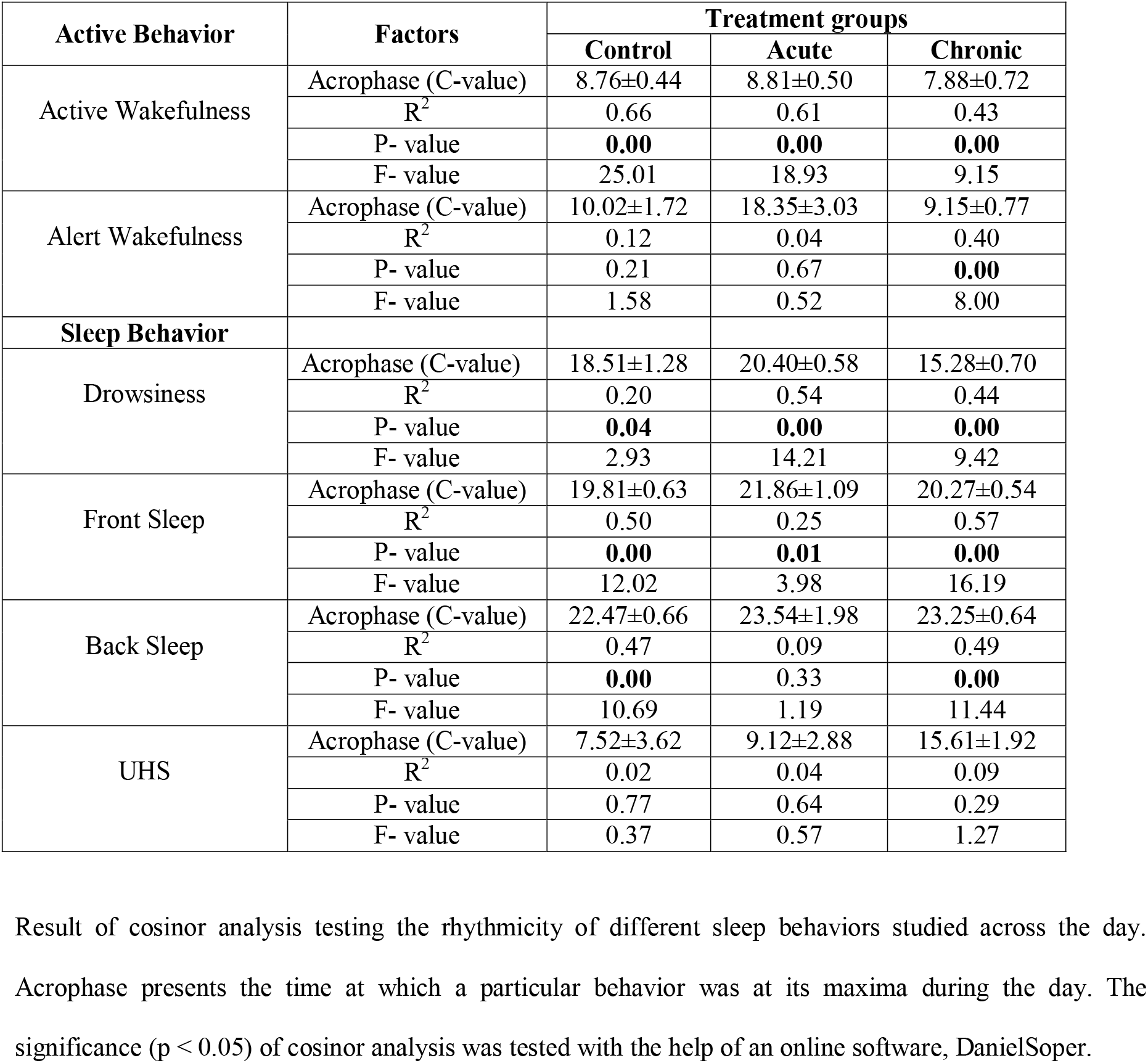
Results of cosinor-analysis

### 3.3.Physiological and metabolic response

Acute exposure of LAN resulted in significant reduction of body mass (P = 0.0007, Student’s t-test; Fig.3a) of the birds under study, despite of having higher food intake (P = 0.0091, Student’s t-test; Fig.3b). Exposure of LAN resulted in temporal alteration of body temperature of birds as well, without affecting its rhythmicity across the day. Chronic exposure of LAN resulted in significant lowering of body temperature of the birds during later half of the night in comparison with acute and control phase of the treatment (interaction; p <.7077, F_10,60_ = 0.714, treatment; P = 0.0234, F_2,60_ = 4.000, time of day; P = 0.0007, F_5,60_ = 5.860; 2-way RM ANOVA, Fig.3c). Measurement of pre and post-prandial glucose reveals the loss of gradient between the two points because of night light (Fig.3d). Maximum prevalence of gradient can be observed in control phase (P = 0.0485, Student’s t-test) which subsequently decreased with acute exposure of LAN (P = 0.1572, Student’s t-test). Chronic LAN exposure results in some sort of rebound in the gradation of blood glucose levels between the two measurement points (P = 0.0316, Student’s t-test). Measurement of plasma corticosterone during midday and midnight points revealed significant effect of LAN on birds’ physiology (Fig.3e). Acute exposure of LAN resulted in significantly higher concentration of plasma corticosterone levels both during midnight and midday in comparison with control and chronic treatments (interaction; P = 0.1950, F_2,16_ = 1.813, treatment; P = 0.0033, F_2,16_ = 8.367, time of day; P = 0.0037 F_1,16_ = 16.44; 2-way RM ANOVA). Besides blood glucose and plasma corticosterone levels, other biochemical parameters such as total protein, creatinine, triglycerides, uric acid, bilirubin total and SGOT were measured from blood plasma (Fig.4). Out of the six biochemical’s mentioned above, level of total protein was affected by LAN both during day (P = 0.043, F_2,8_ = 4.755) and night (P = 0.033, F_2,8_ = 5.353) as revealed by 1-way RM ANOVA (Fig.4a). Diurnal concentration of triglyceride showed elevation in response to acute exposure of LAN (day; P = 0.0004, F_2,8_ = 23.53, night; P = 0.150, F_2,8_ = 2.422) whereas SGOT (day; P = 0.936, F_2,8_ = 0.066, night; P = 0.034, F_2,8_ = 5.278) and creatinine (day; P = 0.518, F_2,8_ = 0.713, night; P = 0.005, F_2,8_ = 10.61) showed variation during the night (Fig.4b, c, d). Uric acid (day; P = 0.330, F_2,8_ = 1.276, night; P = 0.762, F_2,8_ = 0.281) and bilirubin total (day; P = 0.560, F_2,8_ = 0.622, night; P = 0.113, F_2,8_ = 2.890) can be seen affected by night light but greater inter-individual variations resulted in loss of statistical significance of these metabolites (Fig.4e, f).

**Figure 3:**
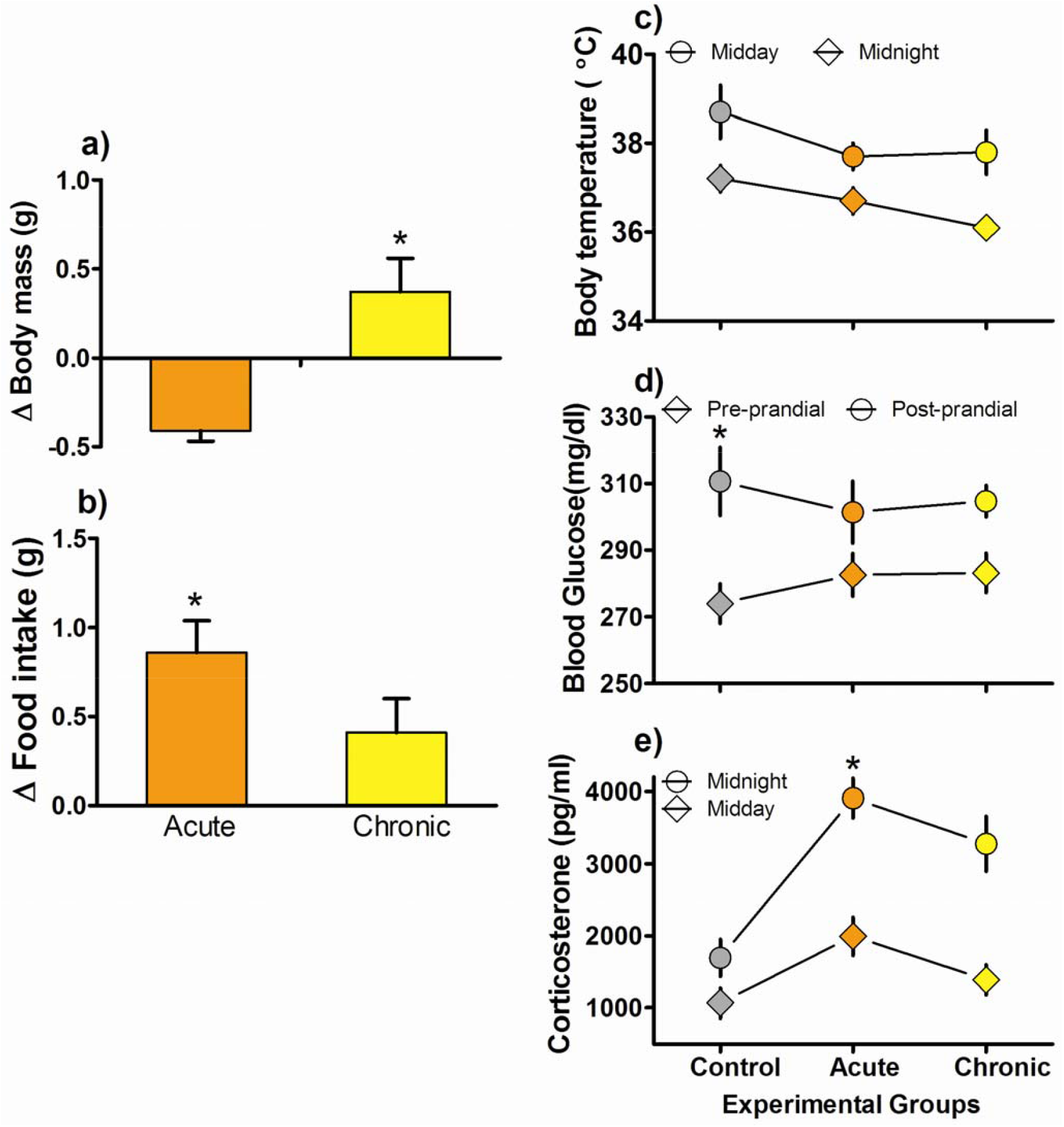
Physiological changes accompanied by LAN exposure. Mean ±SEM values of changes in body mass (a) and food intake (b) along with variation of body temperature at two different time points (ZT-6 and 18) of day (c). Pre- and post- prandial (ZT-2 and 22 respectively) levels of blood glucose was measured against the different treatment schedules (d). Circulating levels of plasma corticosterone was measured during middle of the day (ZT-6) and middle of the night (ZT-18) in each treatment phase during the experiment (e). Asterisk (*) above the bar represent the significant difference between the phases. Significance was considered at p < 00.05.

**Figure 4:**
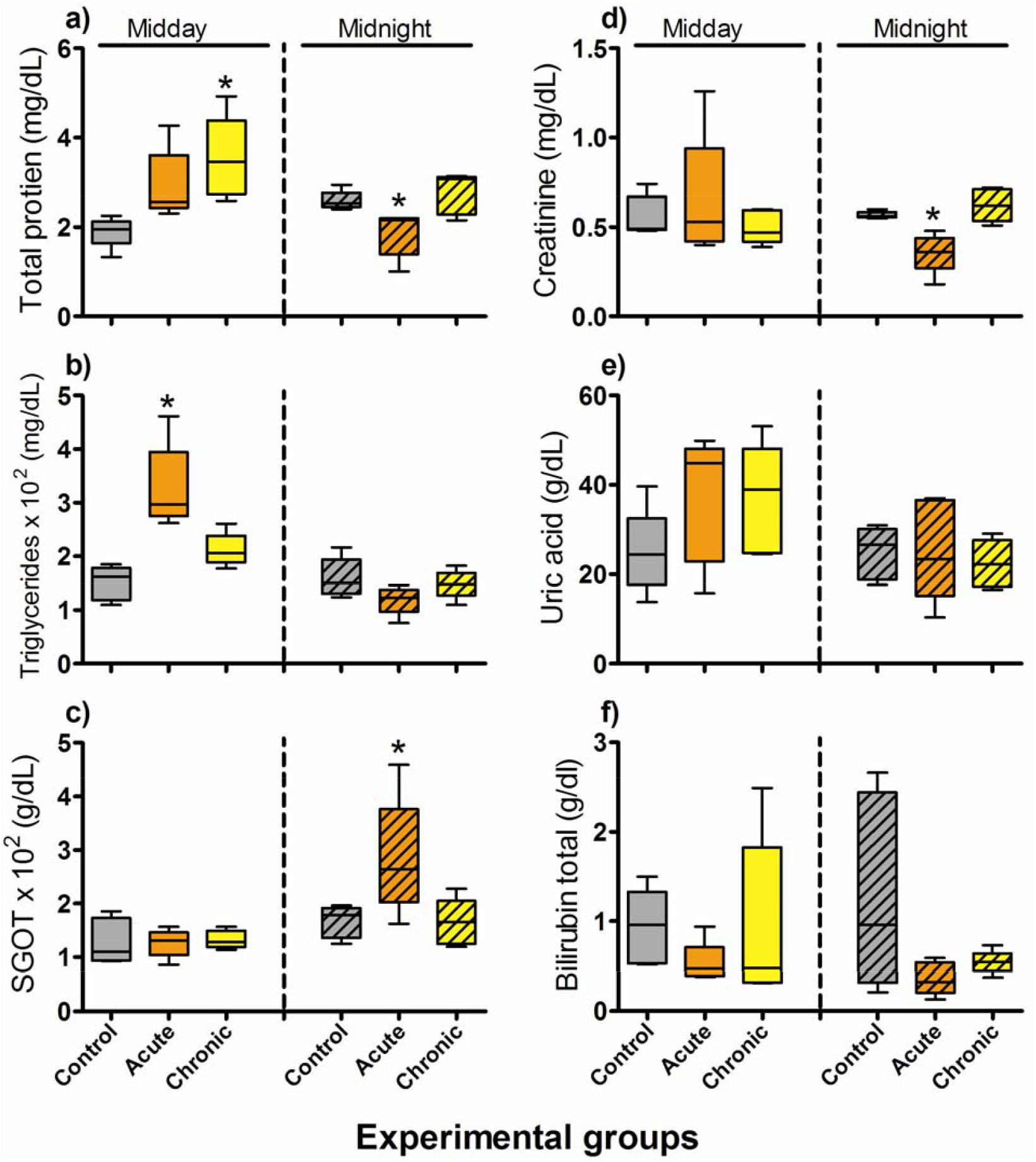
LAN induced daily variation in circulating levels of different plasma metabolites. The concentration of different metabolites is presented as mean ±SEM. The diel variation of total protein (a), triglyceride (b), SGOT (c), creatinine (d), uric acid (e) and bilirubin total (f) at ZT-6 and 18 during different phases of LAN. Asterisk (*) above the bar represent the significant difference between the phases. Significance was considered at p < 00.05.

## 4. Discussion

Our study clearly demonstrates the differential response of different behavioral and physiological parameters with respect to acute and chronic exposure of LAN. In coherence with the studies in the past, our study also reports the prevalence of nighttime disturbance in response to LAN (Ouyang et al. 2017). Acute exposure of LAN resulted in sharp rise of nighttime time activity in a diurnal species ensuing substantial increase in total activity counts of the birds, whereas, chronic exposure of LAN do contributed to the nighttime disturbance, but, with reduced amplitude. Acute LAN exposure drastically altered the activity onset and offset leading to maximum difference between initiation and termination of activity from lights-on and lights-off respectively (de Jong et al. 2017). The initiation of activity advanced by approximately 2.0 hours while the offset delayed by about 1.3 hours. With the passage of time in LAN treatment, the difference between activity onset and lights-on remain nearly fixed indicating towards an irreversible modification of activity onset in response to chronic exposure of LAN. Contrarily the termination of activity gradually came closer towards the actual lights off under the experimental setup. Thus it can be inferred that acute exposure of LAN affects both the initiation and termination of activity but in the long run, it’s the morning activity which is affected by LAN. This can pose survival threat for the birds, as now they can serve as an easy meal for the predators.

As mentioned above, artificial night light, introduced the activity component in the night of a typical diurnal species, thus it has curtailed the share of rest inclusive of its sleep (Raap et al. 2015). Both acute and chronic exposure of night light resulted in significant decrement of nighttime rest when compared with control. Reduced resting duration indicate towards altered sleep behavior. As evident from the results, acute exposure of LAN accounted for 55 percent decrement of sleep whereas chronic exposure of LAN resulted in 14 percent diminution of sleep when compared with control (dark nights). Since sleep constituted of different behavioral postures and was monitored with the help of the same, let’s take a look on the spatial distribution of these behaviors and how they were affected by LAN. As observed in Fig.2a, the different phases of night, namely, early night, mid night and late night showed abundance of front sleep and back sleep postures with minor proportions of drowsiness. With the progression of night, back sleep dominated the bars’. Since, back sleep posture is indicative of deep sleep in birds (Yadav et al. 2021, Ferretti et al. 2019), thus it can be speculated that with the progression of night diurnal birds indulge in deep sleep activity. The transition preceded by late night observation showed 50 percent abundance of front and back sleep postures, while the remaining bar was occupied by drowsiness followed by alert wakefulness. The day component was dominated by active wakefulness, followed by alert wakefulness reporting a typical diurnal attribute. The exposure of LAN has dramatically changed the temporal configuration of different sleep behaviors during the night. Now the different phases of night which were previously occupied by front and back sleep postures were replaced by drowsiness in follow up with alert wakefulness (Fig.2b). The presence of front and back sleep gradually increased with passage of night but could not account for more than 48 percent of the observation period. Thus, behaviorally drowsiness was the significant inclusion in response to introduction of LAN. The possible explanation for this inclusion might be, since drowsiness is connecting link between sleep and activity, thus it offers birds to relax and be vigilant at same time. Not only the night, traces of drowsiness were incorporated even during the day. With the chronic exposure of LAN, the bird sort of, adapted to the circumstances as evident by the relapse of front and back sleep postures in the different phases of night (Fig.2c). The proportion of drowsiness was reduced when compared with acute exposure of LAN, but was not abolished from any phase of nighttime recording. Besides, there was inclusion of drowsiness in every phase of daytime observation suggesting the magnitude of sleep debt in birds because of LAN. Acute exposure of LAN had nearly similar configuration of awake behaviors during the day when compared with control, but, chronic exposure of LAN had significant reduction in active wakefulness. The day was mostly marked by the presence of alert wakefulness followed by drowsiness and UHS. Thus, it can be inferred that long term exposure of LAN results in reduction of active behavior during the day. The plausible reason for reduced locomotion may be sleep debt and energy budgeting (Rezende et al. 2009, Williams and Ternan 1999). Henceforth, it can be speculated that chronic exposure of LAN disrupts the energy balance of the birds.

Besides, reallocation of different sleep and active behaviors, LAN had significant impact on initiation and termination of sleep as well. During dark nights birds had an onset latency of about 20 minutes, which abolished during the acute exposure of LAN. Since we had each observation of 60 minutes, and sleep onset latency was recorded during the transition from day to night, wherein the night component was 30 minute long and we could not find any initiation of sleep in that window for acute exposure of LAN. Chronic LAN resulted in reducing the sleep onset latency thus signifying the sleep debt faced by the birds due to lit-up nights. Moving on to sleep inertia, here too chronic LAN had significantly high values of sleep inertia supporting the preceding statement of sleep debt. Thus long term exposure of night light results in shortening of sleep onset latency and extension of sleep inertia in order to compensate for the sleep loss because of artificial light at night.

Exposure of LAN resulted in higher food intake in birds. Change in food intake with respect to control phase reveals significant higher food intake in acute phase followed by chronic exposure of LAN. Interestingly during acute phase, despite of higher food intake, birds’ had reduced body mass when compared with chronic treatment. This paradox can be answered by significant high levels of nighttime activity, which consumed the excess food intake along with reducing the body mass of the birds. Contrarily, chronic presence of LAN regained the body mass of the birds and reduced the food intake as well, as now the birds had reduced locomotor activity in comparison with acute. Besides affecting the body mass, LAN had significant impact on the body temperature as well. During chronic exposure of LAN, the second half of night specifically reported significantly low body temperatures in birds.

Blood glucose is known to show diel variation (Lobban et al. 2010) with a significant gradient between pre and post prandial glucose levels, but introduction of LAN has removed this gradient between the two points. This response in glucose levels can be attributed to nighttime food intake during acute exposure of LAN, as reported in a recent study from zebra finches (Batra et al.2020). Besides this, advanced activity onset might also be responsible for elevated pre-prandial glucose levels because of night light. With passage of time in LAN, this gradient in glucose between the two points reverted reflecting the adaptability of birds in the prevailing situation. Besides glucose, plasma corticosterone levels also showed the same pattern. Acute exposure of LAN had maximum nighttime corticosterone levels followed by chronic exposure of LAN, which varied significantly when compared with the dark nights. Higher circulating levels of corticosterone can be accredited either to circadian rhythm disruption of HPA axis (Scheving and Pauly 1966; Ouyang et al. 2018) or direct activation of adrenal glands because of night light (Ishida et al. 2005). Higher nighttime activity can also drive higher corticosterone concentration or the reverse can also stand true as reported in some previous studies (Raap et al. 2016a, Alaasam et al. 2018). Besides altered glucose and corticosterone, LAN also affected the circulating levels of total protein and triglycerides in blood plasma of birds. High daytime plasma triglyceride levels can be attributed to higher food intake of birds during acute phase of LAN. Circulating levels of total protein can be seen gradually increasing from acute to chronic treatment during the day. As total protein is sensitive towards levels of dehydration and exposure of LAN resulted in increased activity by the birds, therefore the long term exposure of night light might result in increased levels of dehydration in birds, leading to elevated levels of plasma total proteins. This in turn could result in high viscosity of blood which can lead to cardiac disorders in birds. Concentration of circulating SGOT and creatinine also varied in response to night light. SGOT is a marker for liver damage whereas creatinine represents the muscle breakdown in the body. Since LAN resulted in enhanced activity and sleep loss in birds, this might lead to increased muscle damage and disrupted metabolism in birds. An increasing trend in uric acid concentration can be observed (which corresponds to higher food intake) in the results but because of greater inter-individual differences the significance was lost. Levels of bilirubin total was found unaffected by LAN.

Thus, our study presents an account of behavioral and physiological response of birds against acute and chronic exposure of LAN. As far as locomotor activity and sleep is concerned, on one hand acute treatment results in drastic behavioral alterations but in the long run rebound of these behaviors suggests the adaptive ability of the birds. Nonetheless, the behavioral rebound was accompanied by certain irreversible inclusions such as advanced activity onset and drowsiness. As far as physiology is concerned, relapse in glucose and corticosterone levels can be observed but it does not resemble with that of control, which indicate towards anomalous circadian rhythmicity of the aforesaid parameters in response to long term presence of LAN. Since prevalence of LAN is increasing across the globe and the current study reveals the irreversible alterations in behavior and physiology of birds, such as replacement of sleep posture with drowsiness and elevated levels of certain plasma metabolites and corticosterone, therefore, how these irreversible changes will shape the species survival and existence needs to be taken care of.

## Declarations

## Acknowledgements

The financial support by DBT (Department of Biotechnology) Research Project [BT/PR4984/MED/30/752/2012] to SR and UGC-BSR (Fellowship ID 25-1/2014-2015(BSR)/7-109/2007(BSR) to AY, Department of Zoology, University of Lucknow, Lucknow is highly acknowledged.

## Ethical approval

The experiments were performed as per approval of Institutional Animal Ethics Committee of University of Lucknow, Lucknow India. Protocol number: LU/ZOOL/IAEC/08/17/07 (ii)

## Availability of data and materials

The datasets used and/or analysed in the present study can be obtained from the corresponding author on request.

## Authors’ contributions

Conceptualization and study design: AY, SR and SM; Study conducted by: AY, RK, JT, VV; Statistical analyses: AY; Result interpretation: AY and SR; Manuscript writing: AY and SR. All authors contributed critically to the draft and gave final approval for publication.

## Figure Legends

**Supplementary Figure 1:** Rhythmic profile of different sleep behaviors under LAN. Mean ±SEM values of different sleep behaviors of baya weaver bird during different times of day (ZT-4, 8, 12, 16, 20, 24) and in different treatment phases of experiment.

